# Controlling for variable transposition rate with an age-adjusted site frequency spectrum

**DOI:** 10.1101/2021.08.16.456262

**Authors:** Robert Horvath, Mitra Menon, Michelle Stitzer, Jeffrey Ross-Ibarra

## Abstract

Recognition of the important role of transposable elements (TEs) in eukaryotic genomes quickly led to a burgeoning literature modeling and estimating the effects of selection on TEs. Much of the empirical work on selection has focused on analyzing the site frequency spectrum (SFS) of TEs. But TEs differ from standard evolutionary models in a number of ways that can impact the power and interpretation of the SFS. For example, rather than mutating under a clock-like model, transposition often occurs in bursts which can inflate particular frequency categories compared to expectations under a standard neutral model. If a TE burst has been recent, the excess of low frequency polymorphisms can mimic the effect of purifying selection. Here, we investigate how transposition bursts affect the frequency distribution of TEs and the correlation between age and allele frequency. Using information on the TE age distribution, we propose an age-adjusted site frequency spectrum to compare TEs and neutral polymorphisms to more effectively evaluate whether TEs are under selective constraints. We show that our approach can minimize instances of false inference of selective constraint, but also allows for a correct identification of even weak selection affecting TEs which experienced a transposition burst and is robust to at least simple demographic changes. The results presented here will help researchers working on TEs to more reliably identify the effects of selection on TEs without having to rely on the assumption of a constant transposition rate.

## Introduction

Transposable elements (TEs) were first recognized as important contributors to genetic and phenotypic variation by Barbara McClintock’s pioneering work in maize (1950). Since then, our understanding of the biology and the role of TEs in evolution has increased dramatically (Charlesworth and Charlesworth 1983; Charlesworth and Langley 1986 and 1989; Bennetzen et al. 2005; Tenaillon et al. 2010; Lisch 2013; Barrón et al. 2014; Stuart et al. 2016; Bourque et al. 2018; Bourgeois and Boissinot 2019; Stitzer et al. 2019; Bourgeois et al. 2020) and an increasing number of studies have investigated the genome wide impact of selection on different TE types in various species (e.g. Lockton et al. 2008; Hollister and Gaut 2009; Barrón et al. 2014; Quadrana et al. 2016; Horvath and Slotte 2017; Stritt et al. 2018; Kapun et al. 2020).

Researchers have used a number of approaches to detect selection on TEs. Many have focused on analysis of the site frequency spectrum (SFS), searching for signals that the SFS of TEs deviates from expectations under a particular evolutionary model. However, the transposition rate of TEs is known to be variable and changes in the transposition rate were observed to occur spontaneously or due to stress (Garcia Guerreiro 2012; Belyayev 2014). A non-constant transposition rate is problematic when investigating the SFS and can lead to misleading signals (Bergman and Bensasson 2007; Lockton et al. 2008; Blumenstiel et al. 2014; Bourgeois and Boissinot 2019). For example, a recent rapid increase in the rate of transposition -- a TE burst -- will lead to an overrepresentation of young TEs in the population. Because of their young age, these TEs will predominantly be at a low frequency, mimicking the effect of purifying selection on the shape of the SFS (Bergman and Bensasson 2007; Bourgeois and Boissinot 2019). Note that here we are using the term recent in the sense of evolutionarily recent, not in terms of a few generations ago as is the case in experimental evolutionary studies on TE bursts (e.g. Robillard et al. 2016; Chen et al. 2020). Conversely, if the transposition rate recently decreased, young TEs will be disproportionately underrepresented and skew the SFS towards showing an excess of higher frequency TEs (Bergman and Bensasson 2007; Bourgeois and Boissinot 2019). Thus, shifts in the age distribution of TEs as a result of transposition rate changes can translate into shifts in the SFS that can be misinterpreted as the result of selection acting on the TEs.

Allele frequency is intrinsically linked to allele age. New genetic variants arise in an individual and rise to fixation or become lost from a population over time. Under a neutral infinite sites model in which back mutations do not occur, allele frequency change per generation is both small and stochastic, leading to a positive correlation between allele age and frequency (Maruyama and Kimura 1971; Kimura and Ohta 1973; Maruyama 1974; Watterson 1976; Slatkin 2000; Lynch et al. 2020). This correlation between allele age and frequency can be used to investigate if the alleles are evolving neutrally or under selection (Kimura and Ohta 1973; Maruyama 1974). Alleles which are evolving neutrally will have an allele age-frequency correlation which follows the predictions of Kimura and Ohta (1973). Selection acting on a derived allele changes this relationship, such that, conditional on allele frequency, selected alleles are younger than their neutral counterparts (Maruyama 1974). Previous efforts have capitalized on these predictions to identify selection on individual TE insertions using the mechanics of coalescent theory and a knowledge of population size over time (Blumenstiel et al. 2014).

Because selection changes the age-frequency correlation in predictable directions, analysis of this relationship may provide a useful approach to investigate the evidence of selection on TEs. Thanks to the recent development of new software such as GEVA (Albers and McVean 2020), the age of TEs and putatively neutral sites can be estimated. Hence, the putatively neutral sites can be resampled so that their age distribution matches the age distribution of the TEs. Importantly, comparison of putatively neutral and selected loci stratified to have the same age distribution should resolve a number of issues with inferring selection from the SFS. First, by matching the age distribution of SNPs to that of TEs, differences between SNPs and TEs in the abundance of alleles at given frequencies due to simple neutral processes should be erased. Second, because the comparison is between alleles of similar age distribution, demography will have affected both sets of markers in a similar manner. Finally, many TE datasets depend on ascertaining TEs in one or a few reference individuals. While this skews the frequency distribution, under neutrality matching the age distribution of SNPs and TEs should remove such frequency differences between SNPs and TEs.

In this study, we take advantage of individual based forward-in-time simulations to investigate the utility of comparing the age-frequency distribution of TEs to neutral SNPs. We first demonstrate how a change in transposition rate impacts the allele age-frequency relationship of TEs. We then assess how reliably selection can be detected after conditioning on the age of an allele. Finally, we evaluate how robust this approach is to simple demographic changes such as a population bottleneck and inaccuracies in the age estimates. The results contribute to our understanding of the impact of a non-constant transposition rate on the age and SFS of TEs and showcase how selection on TEs can be investigated without having to assume a constant transposition rate.

## Results

### The Simulated Models

We simulated copy and paste TE polymorphisms under two distinct models using the individual based forward-in-time simulation software SLiM 3.4 (Haller and Messer 2019b) to investigate the correlation between age and frequency of TEs and SNPs in a single population (Figure 1). The first model incorporated a bottleneck that reduced the population size to 10% of its ancestral value (bottleneck model). The second model includes a change in the transposition rate with a 10x increase in the transposition rate of TEs during 250 generations (TE burst model). For both models, five different scenarios with distinct selective constraint on TE insertions were run. The scaled selection coefficients (4*N_e_s*; see methods) used were 0, −0.2, −2, −10, and −20, corresponding to a range from neutral to moderately strong selection against individual TEs. Both models included an ancestral population size of 500 individuals, a 5,000 generation burn-in, and 5,000 generations of evolution following the bottleneck or transposition burst. Each model was run 100 times and sampled at nine different time points to investigate changes in the distribution of allele ages and frequencies.

**Figure 1.**
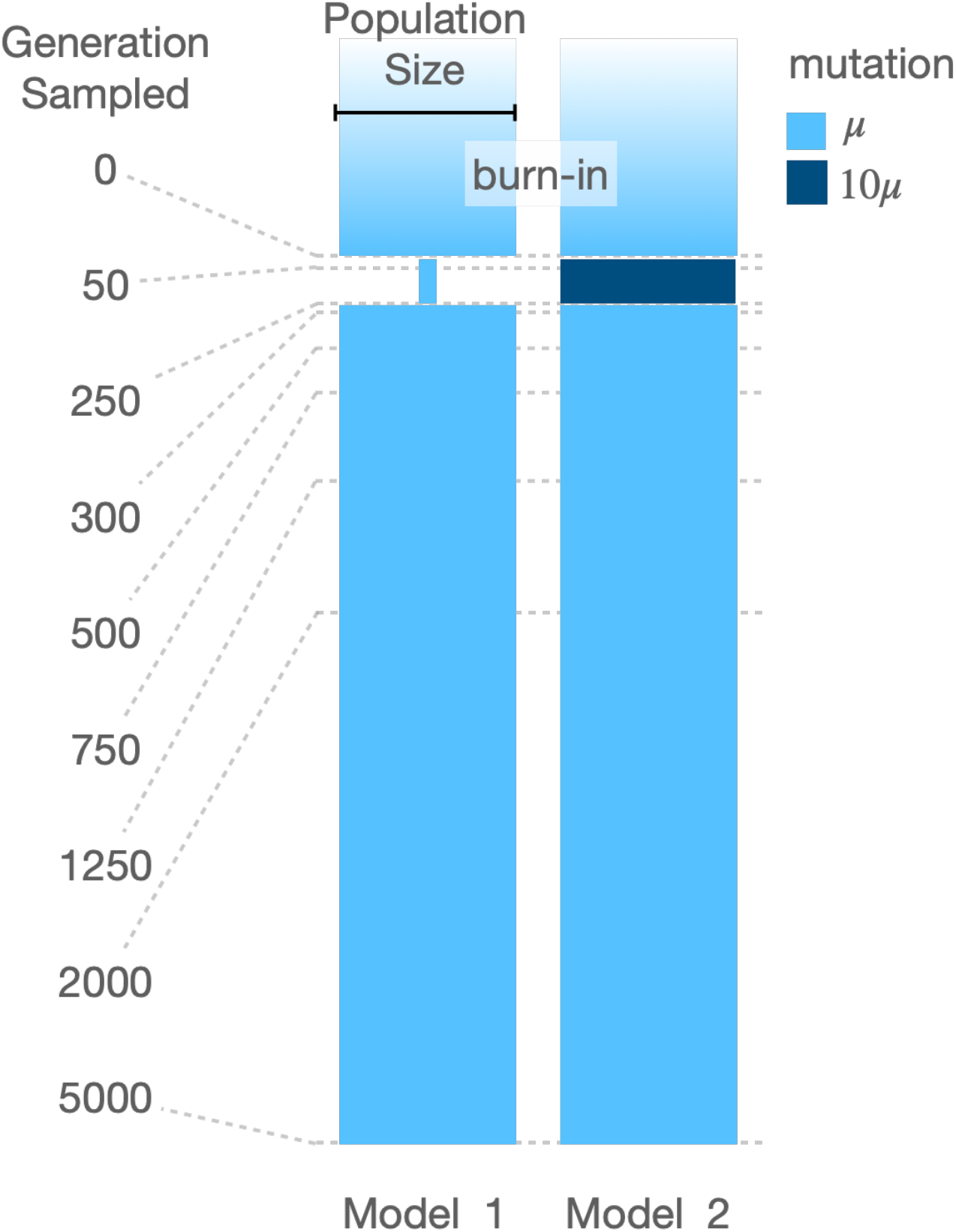
Illustration of the bottleneck (Model 1) and transposition burst (Model 2) models used. The width and height of the blue bars are proportional to population size and time, respectively. The nine sampling time points are represented by dashed lines.

### The Impact of Selection, Demography and Transposition Rate on the Age Distribution

We first investigated the allele age-frequency relationship at the end of the burn-in phase (Generation 0) of the different models. We confirmed that under a constant demography and transposition rate, neutrally evolving TEs and SNPs have an observed mean age distribution, conditional on their allele frequency, which corresponds to the analytical expectations of Kimura and Ohta (1973) (Figure 2A and Supplementary Figure S1A_1_ and S1B_1_). Second, we examined the impact of negative selection on the allele age-frequency relationship of TEs (Figure 2B and Supplementary Figure S1) at Generation 0. We observed that TEs evolving under negative selection are on average younger than neutrally evolving TEs, and the reduction in their age corresponds closely to analytical predictions (Maruyama 1974; Figure 2B and Supplementary Figure S1 and S2). Third, as described by Griffiths and Tavaré (1998), we observe that demographic changes impact the allele age-frequency relationship. Specifically, our demographic model led to a decrease in the mean allele age of neutrally evolving TEs (Figure 2C). The mean allele age decrease observed is less pronounced as the number of generations since the bottleneck increases, indicating that the allele age-frequency distribution is recovering from the impact of the bottleneck through time (Figure 2C). Taken together, the concordance between theory and our simulations indicates that shifts in the allele age-frequency distribution are not due to errors or abnormalities in our simulations but rather these shifts are the results of variations in the parameters used in our models.

**Figure 2.**
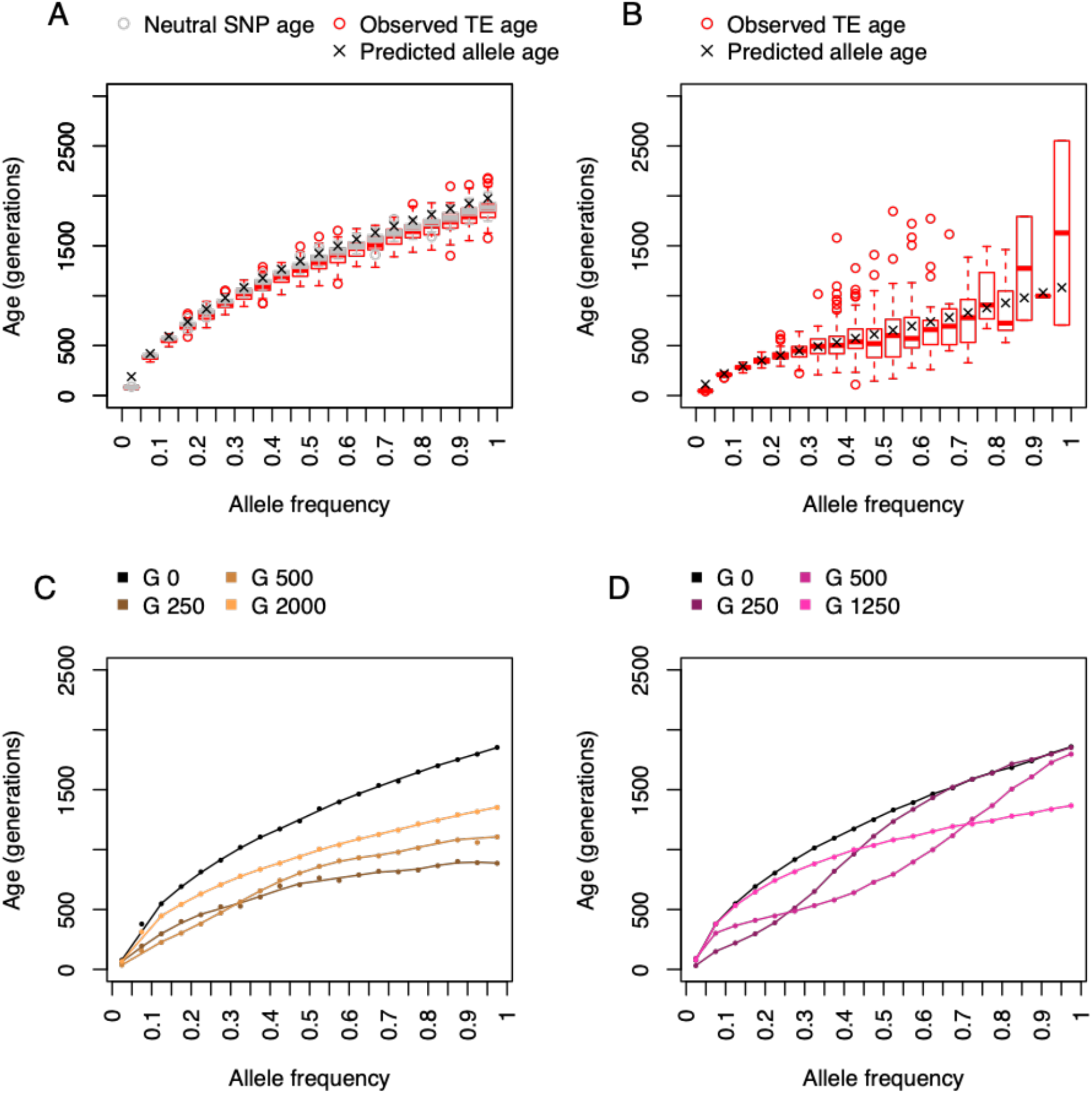
Allele age-frequency relationship. A. Expected (Kimura and Ohta 1973) and observed relationship between mean frequency and allele age of neutrally evolving TEs and SNPs after the burn-in phase (Generation 0). B. Expected (Maruyama 1974) and observed relationship between mean age and allele frequency of TEs under negative selection (4*N_e_s* = −10) after the burn-in phase (Generation 0). C. Effects of a bottleneck on the mean age and allele frequency relationship. D. Effects of a TE burst on the mean age and allele frequency relationship.

Investigating the impact of a TE burst on the relationship between TE age and frequency revealed that indeed transposition rate changes can impact the mean age of TEs at a given frequency (Figure 2D). That the transposition/mutation rate has an impact on the allele age-frequency relationship is not surprising, since Maruyama (1974) showed that the mutation rate impacts the expected age of segregating alleles. The transposition burst simulated here led to a decrease in the mean age of neutral TEs at specific frequencies, depending on the number of generations between the TE burst and the sampling of the population (Figure 2D). If the population was sampled after the transposition rate increased but before it was restored to its starting value (Generation 250 in Figure 2D), TEs at lower frequencies were found to be on average younger than expected but the mean age of high frequency TEs was not affected. However, if the population was sampled after the TE burst ended (Generation 500 and 1250 in Figure 2D), the reduction in the average age of the TEs was observed in TEs with intermediate and higher frequency. This reduction in the average age of TEs caused by the TE burst resembled a wave which passed through the TE frequency distribution through time, leading to a recovery of the allele age-frequency distribution over time similar to the bottleneck model. The wave-like effect of a TE burst on the allele age-frequency relationship indicates that TE insertions which arise before, during or after the TE burst have only a comparatively limited effect on the allele age-frequency relationship and do not alter the entire allele age-frequency distribution as selection or demographic changes would (Figure 2B, C and D). This is similar to the impact of a TE burst on the SFS, where TE insertions which occurred during a TE burst will be disproportionately overrepresented at a specific allele frequency and will, therefore, alter the shape of the TE SFS (Supplementary Figure S3). A disproportionate contribution of TEs with distinct ages to the SFS advocates for a thorough consideration of the age distribution of TEs when investigating frequency shifts observed between TEs and SNPs to avoid false conclusions on the impact of selection on TEs.

### Conditioning on Age to Identify Allele Frequency Shifts Caused by Selection

It has previously been suggested that conditioning on age could help disentangle whether shifts in the population wide frequency distribution of TEs are caused by transposition rate changes or selection (Blumenstiel et al. 2014). To investigate whether choosing neutral alleles that match the age distribution of TEs could enable a reliable identification of shifts in the TE frequency distribution caused by selection, we evaluated our ability to correctly distinguish between TEs evolving under purifying selection and under neutral conditions across our simulations. We resampled SNPs to match the age distribution of the TEs and used an age-binning strategy where the TE and resampled SNP dataset were divided into 10 bins (deciles) based on their age (see Methods). This strategy ensures that the same number of observations were contributing to each bin within a single run and simplifies the comparison between runs simpler as each run had the same number of bins. Using these deciles, we calculated the difference in the mean frequency between TEs and SNPs (Δ frequency: mean TE frequency – mean SNP frequency) for each bin. The observed Δ frequency of the age decile bins were then investigated for a systematic deviation from 0 with increasing age using a rank correlation test (see Methods). Under a strictly theoretical scenario, where we allow our population to have an infinite size and TE are evolving neutrally, we expect the average frequency of TEs within a specific age bin to be equal to the average frequency of SNPs in the same age bin (i.e. Δ frequency = 0). However, in our models Δ frequency can deviate from 0 due to two reasons: genetic drift and/or selection. One key difference between deviations caused by genetic drift and selection is that deviations in Δ frequency caused by selection are directional, which is in contrast to the expected randomness of the deviations caused by genetic drift. Additionally, the magnitude of the deviation of Δ frequency from 0 is expected to be higher in older age bins than in younger age bins, because purifying selection will prevent selected alleles from slowly drifting to intermediate or high frequency and therefore selected alleles can only reach high frequencies through a fast drift to higher frequencies (Maruyama 1974). Additionally, high frequency neutral alleles will be on average older than low frequency neutral alleles (Kimura and Ohta 1973). Therefore, older age bins will include more intermediate and high frequency neutral alleles than young age bins, leading to a higher Δ frequency between selected and neutral alleles in older age bins. Hence, if TEs are under negative selection, we expect Δ frequency to be largest for older bins and a negative correlation between Δ frequency and age.

Using this approach revealed that under ideal conditions of constant demography and transposition rate, fewer than 10% of simulations with neutrally evolving TEs showed a significant correlation between Δ frequency and age. In contrast, in simulations with TEs evolving under purifying selection, a significant negative correlation between Δ frequency and age was observed in at least 95% of the runs (Figure 4 and Supplementary Table S1). Nevertheless, it should be mentioned that our ability to detect a negative correlation between Δ frequency and age was dependent on the selection strength and was severely reduced if the scaled selection coefficient was set to −0.2, corresponding to a nearly neutral fitness effect (Supplementary Table S1). Finding a negative correlation between Δ frequency and age after the burn-in phase in the simulations where negative selection was impacting TEs indicates that, in principle, the approach proposed here can reliably detect the impact of selection on TEs under ideal conditions.

We next explored how demographic changes impact our ability to detect a significant correlation between Δ frequency and age by investigating the Δ frequency and age relationship in our bottleneck model. As in our ideal situation, a significant correlation between Δ frequency and age was rarely observed in our bottleneck simulations, though increased stochasticity due to drift did lead to a much noisier pattern (Figure 3A and 4). However, in runs where TEs were affected by negative selection, the simulated bottleneck limited our ability to detect a significant negative correlation between Δ frequency and age (Figure 3B, 4 and Supplementary Table S1). Such an effect was expected, since demographic changes such as bottlenecks are expected to increase the effect of genetic drift and blur patterns in the frequency distribution caused by selection. Additionally, a reduction in the population size also entails a reduction in the scaled selection coefficient, meaning that selection will be less efficient in removing deleterious TE insertions from the population when the population size is reduced.

**Figure 3.**
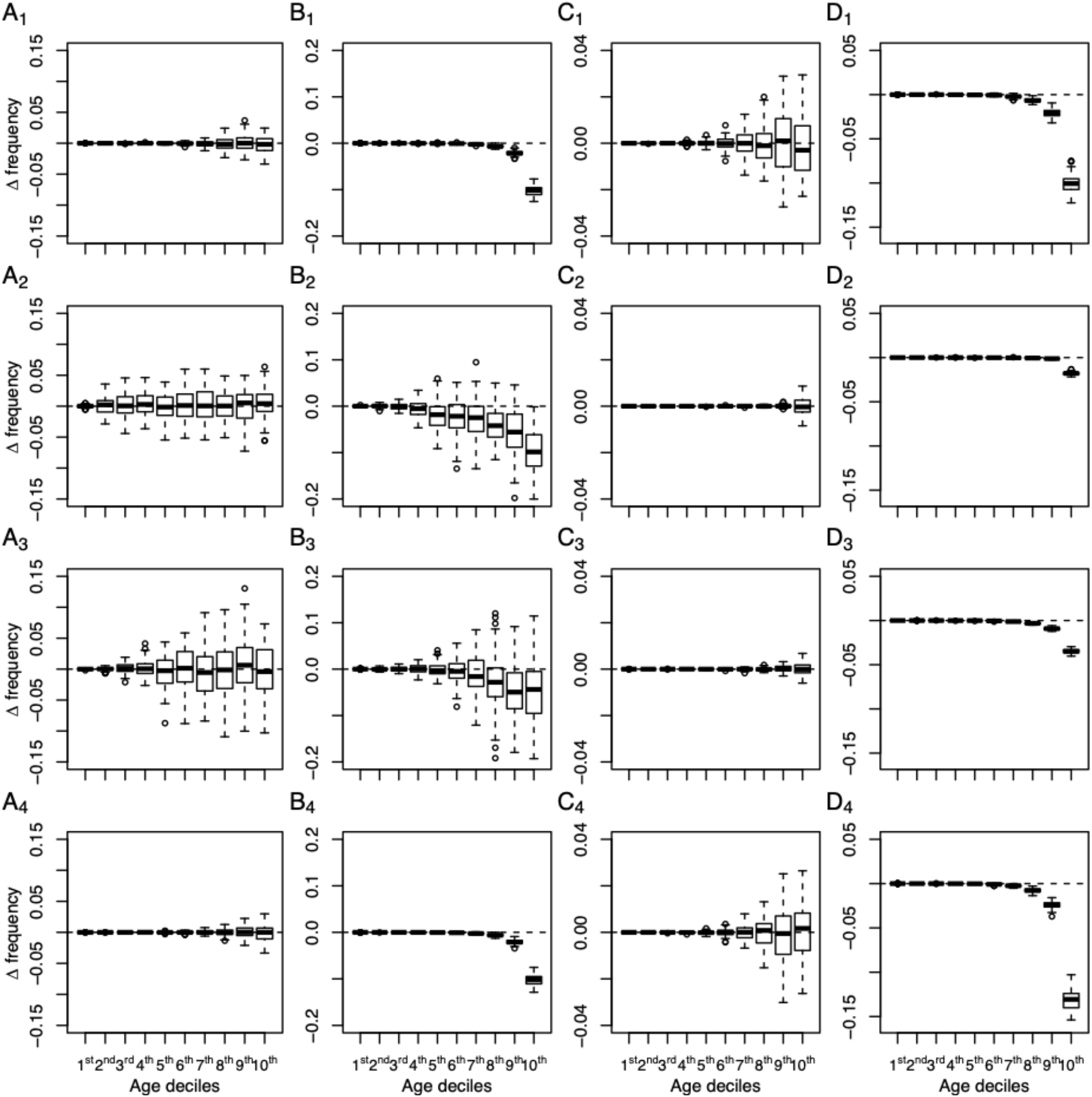
Age binned Δ frequency (mean TE frequency – mean SNP frequency) distributions observed in the different models. A. Observed Δ frequency between neutrally evolving TEs and SNPs under a bottleneck model. B. Observed Δ frequency between negatively selected TEs (4*N_e_s* = −10) and neutrally evolving SNPs under a bottleneck model. C. Observed Δ frequency between neutrally evolving TEs and SNPs under a TE burst model. D. Observed Δ frequency between negatively selected TEs (4*N_e_s* = −10) and neutrally evolving SNPs under a TE burst model. The four rows represent samples from generations 0, 50, 250, and 1250, respectively; generations 300, 500, 2000 and 5000 can be found in Supplementary Figure S4. Note that the y-axis differs across columns.

**Figure 4.**
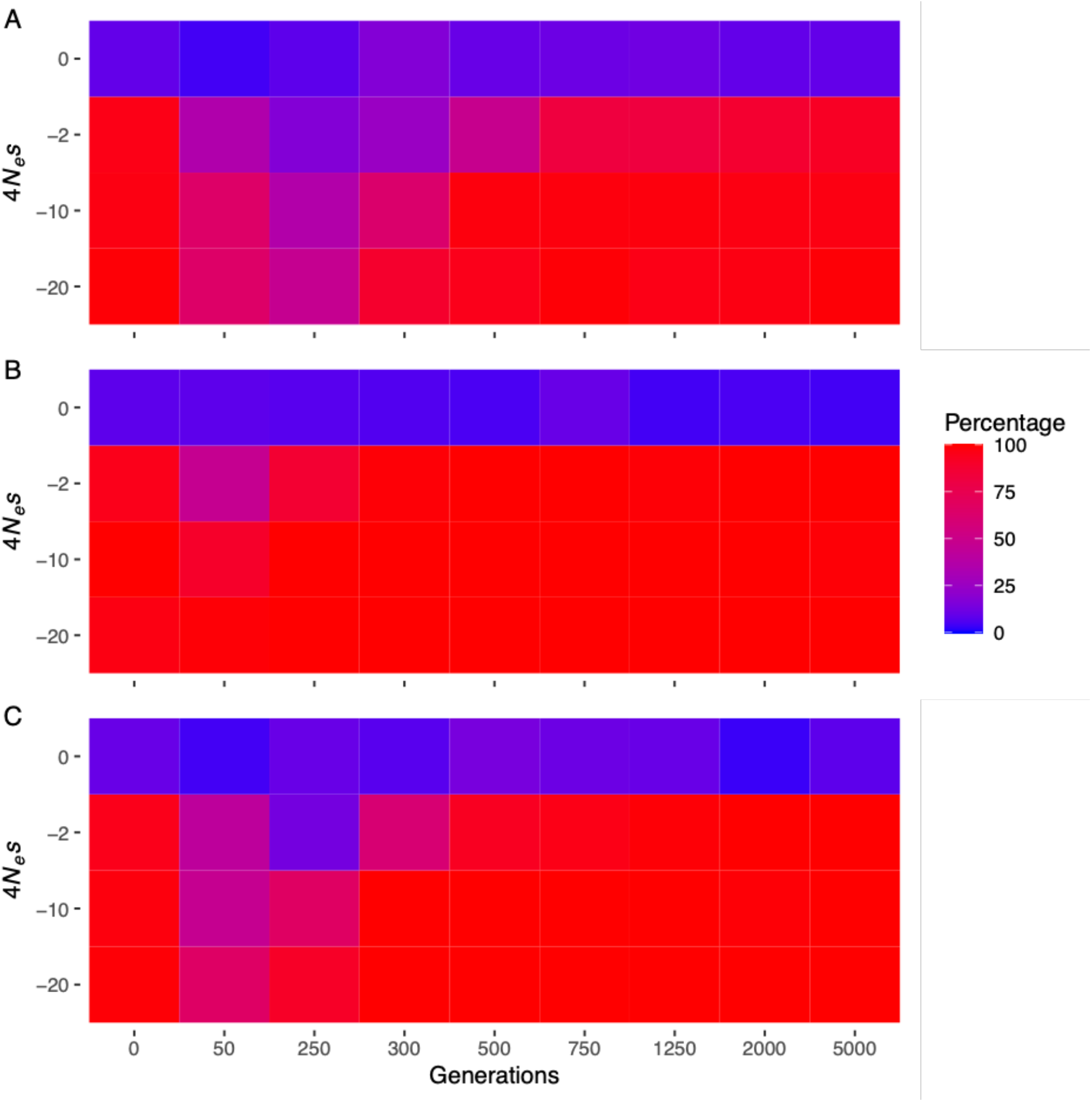
Percent of runs which showed a significant negative correlation between Δ frequency and age at different timepoints (in generations) in our simulations (Spearman’s rank-order correlation test, *p*-value < 0.05). A. Bottleneck model. B. TE burst model. C. Combined bottleneck and TE burst model. The results reported after the burn-in phase at generation 0 correspond to observations under ideal conditions (constant demography and transposition rate). The numerical values underlying the heatmap are given in Supplementary Table S1.

To investigate if conditioning on age was a viable solution to cancel out the effects of a transposition rate change on the frequency distribution of TEs, we investigated the correlation between Δ frequency and age in our TE burst models. Doing so revealed that neutrally evolving TEs had a similar frequency distribution to our neutral sites (Δ frequency scattering around 0; Figure 3C) and in at least 90% of our neutral runs Δ frequency did not deviate from 0 in a systematic way regardless of when the TE burst occurred (Figure 4 and Supplementary Table S1). The false positive rate of our approach when TEs underwent a change in their transposition rate was between 4% and 10%, which was similar to the false positive rate under ideal conditions (Figure 4 and Supplementary Table S1). This contrasts with what is expected when performing a traditional SFS comparison between TEs and SNPs, where a TE burst can lead to shifts in the SFS and to an increased risk of false inferences of selection (Bourgeois and Boissinot 2019; Supplementary Figure S3). In the TE burst model with negative selection, Δ frequency deviated from 0, especially in older site bins (Figure 3D), and a significant negative correlation between Δ frequency and age was observed in more than 95% of our runs (Figure 4). Only when the scaled selection coefficient was set to −2 or less did the TE burst impact our ability to identify a negative correlation between Δ frequency and age (Figure 4 and Supplementary Table S1). After a recent TE burst (generation 50 in our models) our ability to detect a negative correlation between Δ frequency and age was reduced but was quickly restored after selection had time to alter the frequency distribution of TEs which occurred during the burst (Figure 4). Taken together, our simulations show that conditioning on age indeed enables an unbiased comparison of the frequency distribution of TEs to neutrally evolving sites without having to assume a constant transposition rate, and that doing so reduces the risk of falsely inferring selection impacting neutrally evolving TEs while still being able to correctly infer selection on TEs with deleterious fitness effects.

To investigate if our approach could also cope with a model which includes demographic as well as transposition rate changes, we combined our bottleneck and TE burst model to create a model where, after a burn-in phase, a simultaneous decrease in the population size and increase in the transposition rate were modeled. In nature, a strong bottleneck and a TE burst can occur simultaneously. In plants, for example, polyploidization events can be associated with both a strong bottleneck and a TE burst (Vicient and Casacuberta 2017). In the combined model with neutrally evolving TEs, the Δ frequency and age correlation test had a false positive rate between 3% and 14%, similar to the false positive rates observed in each individual model (Figure 4 and Supplementary Table S1). When TEs were under selection, the bottleneck and TE burst mitigated our ability to detect a significant negative correlation between Δ frequency and age, but these effects were also comparable to the observed power losses in the individual models which only included a bottleneck or TE burst (Figure 4 and Supplementary Table S1). These results suggest that potential co-occurrences of demographic changes and a TE burst are not problematic when investigating selection based on an age-adjusted site frequency spectrum analysis.

### Using Empirical Age Estimates

In empirical studies the age of TEs and SNPs is unknown and needs to be estimated, which will inevitably introduce inaccuracies. To evaluate how such inaccuracies impact the performance of our approach, we used Genealogical Estimation of Variant Age (GEVA) (Albers and McVean 2020) to estimate the age of TEs and SNPs in 10 runs of our TE burst and bottleneck model with intermediate selection strength (4*N_e_s* = −10) at generation 300. Singletons were removed from the analysis because GEVA had trouble getting correct age estimates. On average, we obtained age estimates for 80% and 60% of the remaining variants which exhibited a relatively high correlation with true age (Spearman’s rho = 0.69 ±0.0035 and 0.72 ±0.02) for the neutral TE burst and bottleneck model, respectively. Similar deviations of Δ frequency from 0 were observed in the analysis using both estimated and true ages (Supplementary Figure S3B_1_, S3D_1_ and S5). When using the estimated ages, 90% of the runs for the bottleneck model exhibited the expected significant negative correlation between Δ frequency and age while all runs for the TE burst model had a significant negative correlation. These results suggest that our approach can be reliably used even if there are inaccuracies in empirical age estimates, suggesting that our approach is readily applicable to real-world data.

## Discussion

Using simulated data, we showed that adjusting the age distribution of neutral sites to the age distribution of TEs can be used as a reliable approach to distinguish between neutrally evolving TEs and those under selection if the assumption of constant transposition rate cannot be met (Figure 3 and 4). This is crucial, since a non-constant transposition rate can lead to deviations in the site frequency spectrum (Supplementary Figure S3), complicating the evolutionary interpretation of observed TE frequencies (Bergman and Bensasson 2007; Lockton et al. 2008; Blumenstiel et al. 2014; Bourgeois and Boissinot 2019). Here, we highlight that using an age-adjusted site frequency spectrum can reduce the probability of falsely inferring selection on TEs if they are evolving neutrally but with a non-constant transposition rate. Further, we showed that this approach is robust to at least simple demographic changes and inaccuracies in the age estimates. Overall, the approach proposed here to evaluate the observed frequency distribution of TEs is a practical alternative in situations where a constant transposition rate cannot be assumed.

### Detecting Selection Beyond Purifying Selection

This approach is an extension of an SFS comparison between TEs and neutral sites, where shifts in the frequency distribution between TEs and putatively neutral sites are attributed to the effects of selection. Hence, conditioning on age before investigating the SFS of TEs can in principle also be used to detect positive selection, which is expected to lead to a positive correlation between Δ frequency and age. Here we only focused on negative selection, because to be able to detect selection with this approach, selection needs to affect TEs on a larger scale (e.g. genome or TE class/superfamily wide scale) and TEs are in general expected to have a neutral or deleterious fitness effect (e.g. Charlesworth and Charlesworth 1983; Lockton et al. 2008; Barrón et al. 2014). While individual TE insertions have been shown to be adaptive (González et al. 2008 and 2010; Rishishwar et al. 2018; Bourgeois and Boissinot 2019), a wide scale analysis of the TE frequency distribution will miss such instances. However, TE insertions under longstanding balancing selection will stand out during the process of age conditioning, since such TE insertions are expected to be older than other sites and finding segregating neutral sites of approximately the same age will be challenging. Such TE insertions might need to be excluded before investigating the correlation between Δ frequency and age. Hence, investigating the age of TEs can lead to the identification of candidate insertions under balancing selection if other causes of old segregating TEs, such as TE excision after reaching fixation (see below), can be excluded.

### The Impact of TE Biology

While we tried to make realistic models to simulate the evolution of TEs, TE biology is much more complex than modeled here. Some aspects of our simulations, such as the possibility of TEs becoming inactive (see methods), were chosen to represent some of the complex biology of TEs, but several simplifications have been made which might need to be considered when applying this approach on real data.

In our models, we limited the transposition mechanism of our TEs to only a copy and paste transposition mechanism. A cut and paste transposition mechanism, which can cause a TE to exit an insertion site, was not modeled in order to avoid the need of considering back mutations or a third allele at insertion sites (if alleles which never had a TE insertion are seen as different from alleles were the TE was cut out due to processes like imprecise excision (Hickman and Dyda 2016)). This simplification allowed us to use the work from Kimura and Ohta (1973; eq 13) and Maruyama (1974; eq 13), which were derived for biallelic sites with no back mutations, to predict the expected allele age-frequency relationship for the different simulations. However, if TEs can be excised from an insertion site, this can lead to a decrease in the population wide allele frequency of the TE which is not due to selection, and such back mutations could lead to shifts in the allele age-frequency relationship (Maruyama 1974). For example, a TE could reach fixation and afterward be cut out and lead to a segregating TE insertion which is older than expected based on its frequency.

Another simplification made in our models was to assume that active and disabled TEs have the same fitness effect. This assumption was done in order to be able to consider sites with active, disabled and no TE insertions in the population as biallelic loci (one allele with the TE insertion, regardless of its activity status, and one without insertion). This simplification was again made to be able to use the theoretical results derived for biallelic loci from Kimura and Ohta (1973) and Maruyama (1974). Nevertheless, it should be noted that active and disabled TEs can have distinct fitness effects, for example disabling a TE through epigenetic silencing can decrease the fitness of the host if nearby functional sequences are accidentally silenced along with the targeted TE (Hollister and Gaut 2009). Further, we should also note that it is known that TEs come with many different characteristics and insertion preferences (Bourque et al. 2018) which were ignored in our models. As the main goal of this study was to show the principle that conditioning on age is a viable way to investigate selection on TEs, we argue that the simplifications made here are justifiable. Using this approach on real data will, however, require an individual evaluation of the characteristics of the investigated TEs and their potential impact on the expected correlation between Δ frequency and age. In cases where TE groups may differ in the Δ frequency and age correlation, splitting up the TE dataset based on, for example, TE class/superfamily, genomic regions, epigenetic status or other criteria, or the exclusion of some TEs before running the selection analysis should be considered.

### Limiting Factors

As with any method, our approach comes with some limitations, besides the obvious requirement of being able to estimate the age of the TEs and the neutral sites. First, selection on TEs needs to be strong enough to be able to detect shifts in the frequency distribution caused by selection. The stronger the selection strength against TE insertions the better our ability to detect a significant negative correlation between Δ frequency and age (Figure 4 and Supplementary Table S1). If the scaled selection coefficient was set to −0.2, which corresponds to a nearly neutral fitness effect, our approach had difficulty detecting frequency shifts caused by selection (Supplementary Table S1). However, for a scaled selection coefficient of −2, which is usually associated with quite weak selection, we were still able to detect negative correlations between Δ frequency and age caused by selection. Hence, selection against TEs does not need to be strong to be detected with this method. At the other extreme, very strong scaled selection coefficient (< −100) may also cause problems when trying to identify a negative correlation between Δ frequency and age because in such cases all segregating TE insertion are expected to be quickly removed by selection and, therefore, most TE insertions observed in the population will be very young with only a few older segregating TE insertions (results not shown). A small number of old TEs will decrease our ability to observe a negative correlation between Δ frequency and age. The number of segregating TEs in the analyzed dataset is a second limiting factor, since fewer datapoints will lead to more noise and a reduction in the power of the analyzes. In our simulation, the number of segregating TEs varied over a couple of magnitudes between bottleneck and TE burst models (Supplementary Table S2), which contributed to our ability to detect the effects of negative selection at the different time points. But, as a point of reference after the burn-in phase we had between 3,000 and 10,000 segregating TEs (Supplementary Table S2).

Finally, when comparing TEs to putatively neutral sites, the putatively neutral sites may not be evolving neutrally, either because of direct or linked selection. It has been shown that even synonymous sites can be under selective constraint (Chamary and Hurst 2005; Chamary et al. 2006; Eöry et al. 2010; Künstner et al. 2011; Gu et al. 2012; Lawrie et al. 2013; Gossmann et al. 2018) and, therefore, the SFS of putatively neutral sites such as synonymous sites might be altered by natural selection. However, having weak selection affecting the putatively neutral sites might not be as important as the difference in the strength of selection between the compared sites, since the approach proposed here is evaluating frequency differences between sites of the same age. Hence, if selection against TEs is strong enough, a correlation between Δ frequency and age should be detectable even if the putatively neutral sites are under weak selective constraint. In our simulations neutral sites were not affected by direct selection, but some effects of linked selection on the neutral sites especially during a TE burst could not be completely excluded. In order to minimize the impact of linked selection, the recombination rate was set high enough to ensure that after the burn-in phase the SFS of our neutral sites was not altered by linked selection, which was checked by comparing the SFS of SNPs between the different models with varying selection strength (Supplementary Figure S6). Here, we were able to detect a negative correlation between Δ frequency and age even in our TE burst model with the strongest selective constraint against TEs, where linked selection on neutral sites is expected to be the strongest.

## Conclusion

When investigating the SFS of TEs for signs of selective constraints, it is important to consider if a constant transposition rate can be assumed. If such assumption cannot be made, using an age-adjusted site frequency spectrum when comparing TEs to neutral sites can reduce the risk of falsely inferring a selective constraint on neutrally evolving TEs, where shifts in the SFS were caused by changes in the transposition rate. Here, we presented one way to condition on age before comparing the allele frequency of TEs to the allele frequency of neutral sites, and showed that our approach can reliably detect shifts in the SFS caused by selection even after demographic and transposition rate changes occurred. Further, a change in the transposition rate did not increase the false positive rate of inferring selection on neutrally evolving TEs when using an age-adjusted SFS comparison. While this approach has some caveats, we argue that thanks to new methods to estimate the age of TEs and SNPs, conditioning on age when investigating selection on TEs is a reliable way to account for non-constant transposition rates.

## Materials and Methods

### Model Simulations

In this study, we performed individual based forward-in-time simulations using SLiM 3.4 (Haller and Messer 2019a, 2019b) to simulate the evolution of TEs under different conditions. SNPs and TEs were simulated following the recipe 13.16 “Modeling transposable elements” provided in SLiMgui (Haller and Messer 2019a, 2019b), which was adjusted to reflect the demographic and transposition rate changes investigated. In all models, we simulated a 3Mb genomic fragment in a population of 500 diploid individuals with a nucleotide mutation rate of 10^−6^ mutation/site/generation. Recombination rate was set at 10^−4^/site/generation for the first and last 500kb of the genomic fragment, 5×10^−5^/site/generation for the second and second to last 500kb of the genomic fragment, and at 10^−5^/site/generation for the middle 1Mb of the genomic fragment to mimic the structure of a typical metacentric plant chromosome. Recombination rate was set high enough to ensure linked selection did not impact our neutral sites, which we checked by investigating the SNP SFS of the different models after the burn-in phase (Supplementary Figure S6).

TEs were allowed to evolve under five different selective constraints by setting a constant selection coefficient in SLiM such that the scaled selection coefficient (S) corresponded to: 0 (a neutral fitness effect); −0.2 (a nearly neutral fitness effect); −2 (a weak deleterious fitness effect); −10 (an intermediately deleterious fitness effect) and −20 (a strong deleterious fitness effect). Here S was defined as 4*N_e_s* following Tataru et al. (2017), with *s* being the strength of selection affecting a heterozygous individual assuming no dominance effects. Note that the fitness of homozygous derived genotypes was defined as 1-2s in the work of Maruyama (1974) but as 1-*s* in SLiM. TEs had a transposition probability of 0.0001/copy/generation following a copy and paste transposition mechanism and each individual TE copy had a probability of 5×10^−5^ to be disabled per generation. Disabling an active TE reduces its probability of being copied to 0 but does not alter its selection coefficient.

Each simulation was started by introducing 5000 fixed TEs at random positions in the genome of our individuals, which was then followed by a burn-in phase of 5000 generations where SNPs and TE insertions were allowed to occur and accumulate according to their respective selective constraints. After the burn-in phase, a second phase of 5000 generations started where the different demographic and transposition rate changes were modelled (Figure 1). Our bottleneck model (model 1) consisted of a reduction in the population size to 50 individuals from generation 1 to 250 after the burn-in phase. The TE burst model (model 2) consisted of a ten-foldincrease of the transposition rate to 0.001/copy/generation from generation 1 to 250 after the burn-in phase. Finally, a model which included a demographic and transposition rate change was made by combining the bottleneck and TE burst models, where a reduction in the population size to 50 individuals and an increase of the transposition rate to 0.001/copy/generation occurred from generation 1 to 250 after the burn-in phase. Each model was run 100 times to investigate variation within each model.

From each model we sampled the entire population at multiple timepoints (Figure 1). The position, age and frequency of each SNP and TE insertion was recorded for each timepoint and saved in a datafile using built-in SLiM commands.

### Conditioning on Allele Age and Allele Frequency Analysis

To investigate the allele age-frequency relationship, the output files from our simulations were analyzed in R (R Core Team 2019) using custom scripts (https://github.com/Roberthorv/age-adjusted_site_frequency_spectrum). To contrast our observations with the theoretical results from Kimura and Ohta (1973) and Maruyama (1974), we implemented these results in R (R Core Team 2019).

To compare TE and SNP data with an identical age distribution, we resampled our SNPs without replacement to exactly match the age distribution of the segregating TEs in each run separately. If fewer SNPs were observed within a specific age than TEs, we resampled the SNP dataset with replacement. If no SNPs were found to have the exact same age as our TEs, we increased the age interval of our target SNPs by plus and minus one generation until some SNP datapoints were available and resampled this dataset to include exactly the same number of datapoints as the TE dataset. Once the SNP resampling was done, both the TE and resampled SNP data were binned into 10 age bins each containing 10% of the datapoints (deciles). The age decile bins were then used to calculate the difference in the mean frequency between TEs and SNPs (Δ frequency: mean TE frequency – mean SNP frequency) per bin. To test for signs of selection altering the frequency distribution of TEs, Δ frequency was investigated for a systematic deviation from 0 by performing a one-sided Spearman’s rank correlation test between age bin and Δ frequency.

### Inferring Allele Ages

To evaluate the efficacy of our age-adjusted site frequency spectrum approach to detect selection on TEs when true ages are unknown, as in empirical datasets, we inferred allele ages using a non-parametric approach implemented within Genealogical Estimation of Variant Age (GEVA) (Albers and McVean 2020). GEVA relies on pairwise differences in identity by descent (IBD) regions around the focal mutation to estimate a posterior distribution for the time of origin of the mutation and thus can be implemented for any mutation type. We estimated the TE and SNP ages in our simulations by sampling the genotypes of all our individuals in the TE burst (model 1) and bottleneck (model 2) model with intermediate selection strength (4*N_e_s* = −10) at generation 300. For each model we selected 10 random runs and used a custom R script to prepare the output vcf files from SLiM for an analysis with GEVA (https://github.com/Roberthorv/age-adjusted_site_frequency_spectrum). While GEVA is largely unbiased by the underlying demographic history, we simulated 1000 haploid individuals (up to 5 million IBD segments) for each model under neutrality in msprime v.1.0 (Kelleher et al. 2016) to generate the emission and initial state probability matrices that are used as inputs for inferring ages (https://github.com/Roberthorv/age-adjusted_site_frequency_spectrum). GEVA was run using a genetic map generated based on the parameters used in our simulations (see above Model Simulations section). Similar to the conditioning on age approach used for true ages, we matched the age distribution of SNPs to mirror the one of TEs, binned the data in deciles, calculated Δ frequency for each bin and identified systematic deviation from zero in older age bins.

## Acknowledgments

The authors thank the whole Ross-Ibarra lab for helpful discussions. This work was supported by the National Science Foundation division of Integrative Organismal Systems grant number 1934384 and 1907343. We would like to acknowledge Felix Andrews for the statistical advice, although we did not follow it.

## Data Availability

The SLiM, R, msprime and GEVA scripts used in this study are available on github: https://github.com/Roberthorv/age-adjusted_site_frequency_spectrum

