## Supplementary material for "Controlling for variable transposition rate with an age-adjusted site frequency spectrum": Horvath et al. Supplementary Material.pdf

**Supplementary Tables**

**Table S1**. Percent of runs which showed a significant negative correlation between Δ frequency and age at different timepoints (in generations: G) in our simulations ($p$-value < 0.05). The third column indicates the observations made after the burn in phase at generation 0 (G 0), which corresponds to ideal conditions (constant demography and transposition rate).

| Model | $4*N_e*s$ | G 0 | G 50 | G 250 | G 300 | G 500 | G 750 | G 1250 | G 2000 | G 5000 |
|---|---|---|---|---|---|---|---|---|---|---|
| Bottleneck | | | | | | | | | | |
| | 0 | 9% | 4% | 8% | 17% | 10% | 11% | 12% | 9% | 9% |
| | -0.2 | 26% | 7% | 6% | 16% | 13% | 16% | 20% | 28% | 32% |
| | -2 | 96% | 35% | 17% | 25% | 48% | 82% | 82% | 88% | 92% |
| | -10 | 97% | 61% | 31% | 69% | 94% | 97% | 98% | 98% | 99% |
| | -20 | 99% | 65% | 47% | 89% | 95% | 99% | 96% | 97% | 99% |
| TE burst | | | | | | | | | | |
| | 0 | 8% | 8% | 7% | 6% | 5% | 10% | 4% | 5% | 4% |
| | -0.2 | 29% | 10% | 27% | 15% | 28% | 35% | 44% | 45% | 48% |
| | -2 | 95% | 47% | 87% | 99% | 100% | 100% | 99% | 100% | 100% |
| | -10 | 98% | 90% | 100% | 100% | 100% | 100% | 100% | 100% | 100% |
| | -20 | 97% | 99% | 100% | 100% | 100% | 100% | 100% | 100% | 100% |
| Bottleneck and TE burst | | | | | | | | | | |
| | 0 | 10% | 4% | 10% | 7% | 14% | 11% | 10% | 3% | 8% |
| | -0.2 | 23% | 10% | 5% | 9% | 26% | 28% | 27% | 38% | 44% |
| | -2 | 95% | 42% | 13% | 60% | 93% | 96% | 99% | 100% | 100% |
| | -10 | 98% | 47% | 67% | 100% | 100% | 100% | 100% | 99% | 100% |
| | -20 | 99% | 66% | 91% | 100% | 100% | 100% | 100% | 100% | 100% |

**Table S2**. Range (minimum – maximum) of segregating TE insertions in the population at different timepoints (in generations: G) in our simulations.

| Model | 4*$N_e$*$s$ | G 0 | G 50 | G 250 | G 300 | G 500 | G 750 | G 1250 | G 2000 | G 5000 |
|---|---|---|---|---|---|---|---|---|---|---|
| Bottleneck | | | | | | | | | | |
| | 0 | 9 238-9 853 | 2 518-2 830 | 856-1 018 | 4 937-5 413 | 6 839-7 407 | 7 713-8 298 | 8 684-9 445 | 9 590-10 403 | 11 773-12 643 |
| | -0.2 | 8 724-9 447 | 2 388-2 753 | 793-985 | 4 748-5 218 | 6 569-7 084 | 7 295-7 951 | 8 202-9 008 | 9 025-9 871 | 10 662-11 716 |
| | -2 | 6 183-6 654 | 1 402-1 665 | 563-728 | 3 567-3 945 | 4 854-5 272 | 5 335-5 847 | 5 809-6 322 | 6 086-6 624 | 5 935-6 523 |
| | -10 | 3 656-4 035 | 489-628 | 346-449 | 2 746-3 052 | 3 453-3 812 | 3 564-3 932 | 3 420-3 836 | 3 349-3 704 | 2 893-3 213 |
| | -20 | 2 969-3 349 | 327-455 | 283-396 | 2 551-2 829 | 2 945-3 277 | 2 875-3 239 | 2 884-3 156 | 2 746-3 040 | 2 365-2 571 |
| TE burst | | | | | | | | | | |
| | 0 | 8 987-9 846 | 53 024-57 161 | 87 204-93 648 | 42 606-45 500 | 30 185-32 445 | 28 242-30 116 | 28 449-30 988 | 30 717-33 119 | 46 721-50 537 |
| | -0.2 | 8 604-9 410 | 51 400-53 960 | 83 653-88 956 | 40 684-43 422 | 28 466-30 442 | 26 504-28 315 | 26 698-28 499 | 28 124-30 239 | 40 560-44 221 |
| | -2 | 6 191-6 664 | 39 188-41 480 | 62 226-65 809 | 29 189-30 879 | 19 538-20 764 | 17 085-18 216 | 15 804-16 899 | 15 322-16 555 | 15 748-17 233 |
| | -10 | 3 656-4 004 | 29 406-31 292 | 39 631-42 590 | 16 049-17 397 | 9 451-10 204 | 7 915-8 533 | 7 172-7 711 | 6 818-7 332 | 5 894-6 357 |
| | -20 | 3 030-3 317 | 26 411-28 279 | 31 355-33 494 | 10 588-11 675 | 6 477-7 034 | 5 883-6 443 | 5 694-6 226 | 5 485-5 992 | 4 678-5 225 |
| Bottleneck and TE burst | | | | | | | | | | |
| | 0 | 9 209-9 961 | 6 851-7 576 | 8 568-9 657 | 17 570-19 128 | 20 459-22 034 | 22 242-23 953 | 25 377-27 179 | 29 474-31 339 | 47 012-50 646 |
| | -0.2 | 8 718-9 416 | 6 728-7 369 | 8 216-9 283 | 16 866-18 206 | 19 552-21 129 | 21 427-23 072 | 24 048-25 673 | 27 444-29 238 | 41 693-44 787 |
| | -2 | 6 206-6 714 | 4 743-5 311 | 6 215-7 047 | 12 709-13 694 | 14 322-15 692 | 15 251-16 404 | 15 913-17 110 | 16 363-17 656 | 17 650-19 046 |
| | -10 | 3 653-3 998 | 2 982-3 458 | 4 207-4 954 | 8 690-9 737 | 8 959-9 739 | 8 491-9 120 | 7 829-8 439 | 7 358-8 000 | 6 333-6 923 |
| | -20 | 3 012-3 320 | 2 606-3 134 | 3 376-4 013 | 7 222-7 875 | 6 769-7 254 | 6 359-6 822 | 5 994-6 517 | 5 766-6 361 | 4 960-5 479 |

**Supplementary Figures**

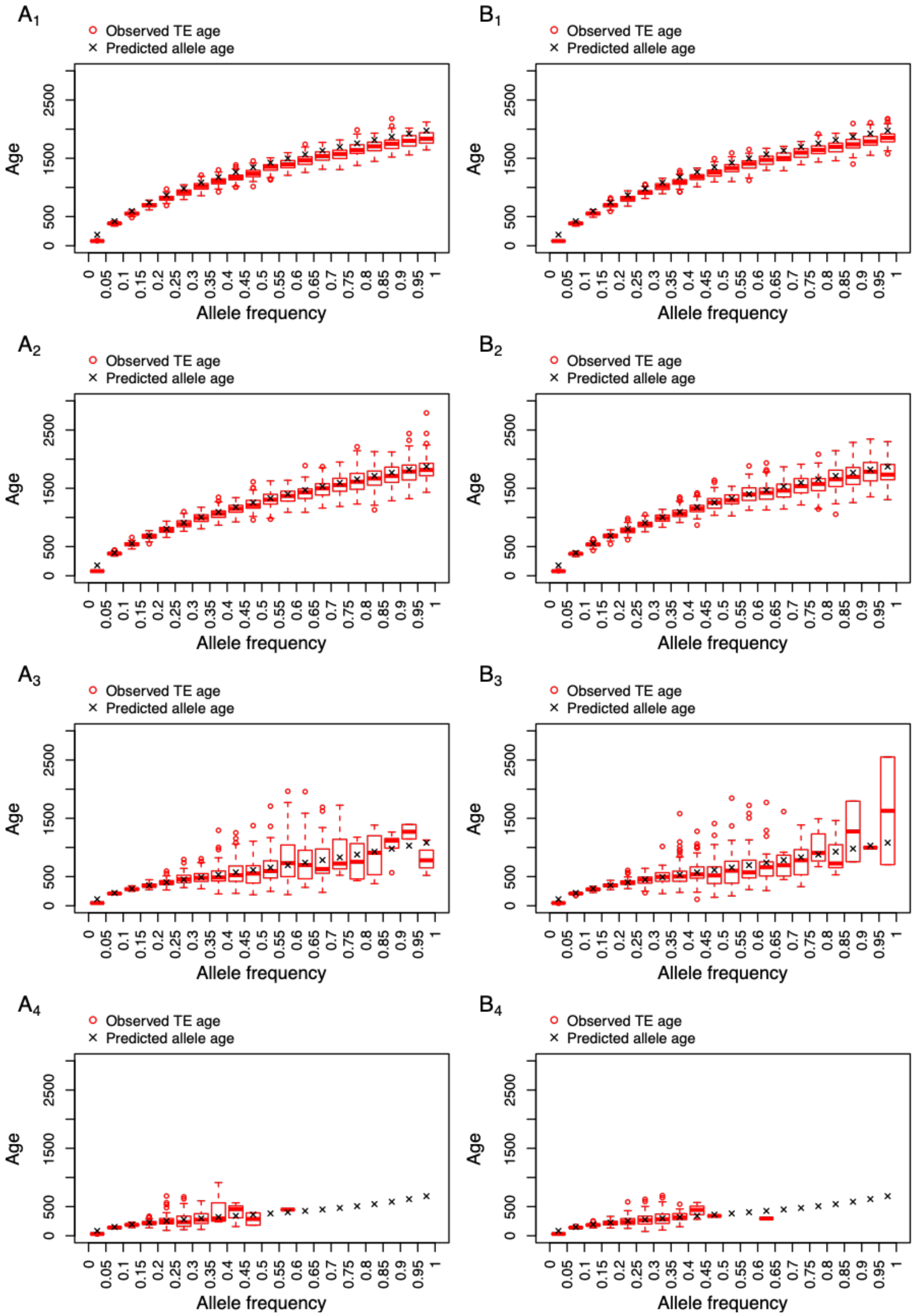


**Figure S1.** Expected (Kimura and Ohta 1973; Maruyama 1974) and observed mean age distribution of TEs at a specific frequency based on 100 runs after the burn in phase (Generation 0). Plot $A_1$ – $A_4$: bottleneck model and plot $B_1$ – $B_4$: TE burst model. The subscript 1 to 4 indicates the strength of selection with the scaled selection coefficient corresponding to 0, -2, -10, -20, respectively.

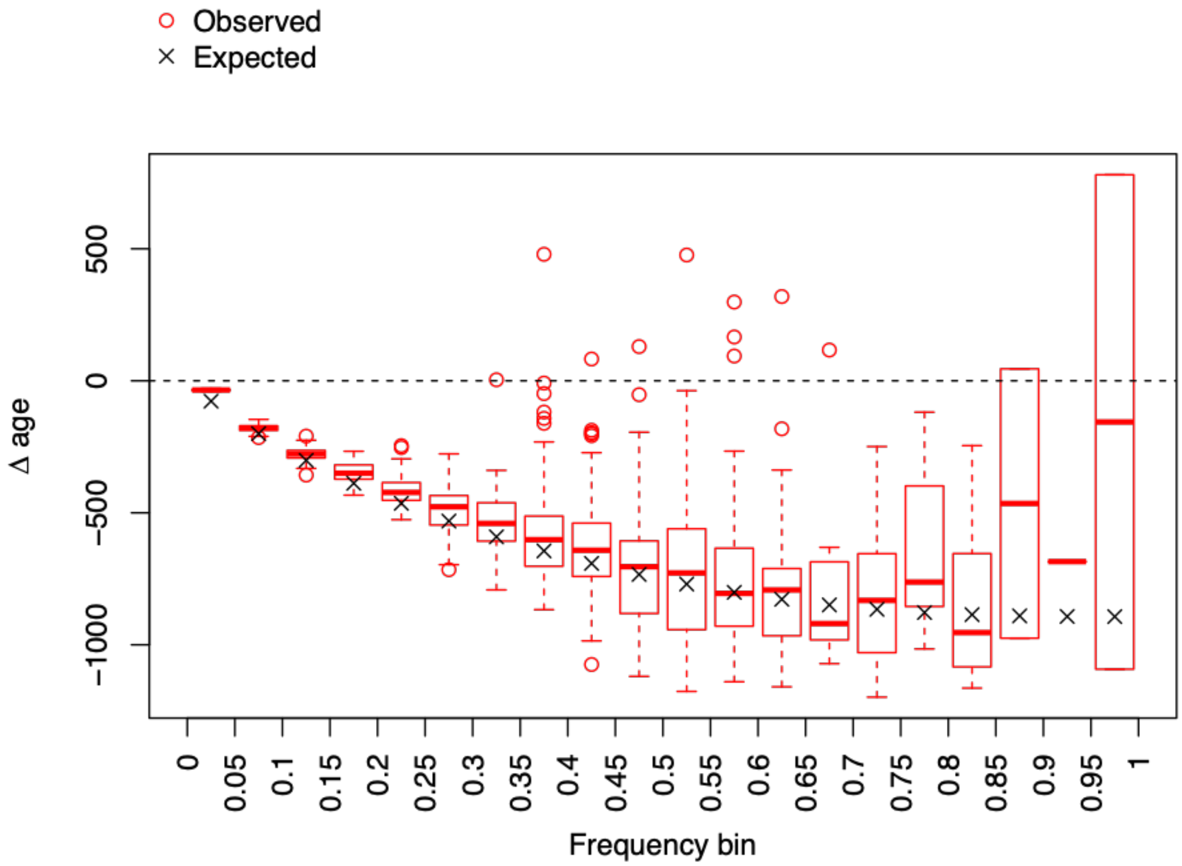


**Figure S2**. Difference in age between TEs and SNPs (Δ age: mean TE age – mean SNP age) at a specific frequency caused by negative selection. Observed Δ age between TEs under selection (scaled selection coefficient -10) and neutrally evolving SNPs are shown in red and expected Δ age (mean allele age predicted by Maruyama (1974) - mean allele age predicted by Kimura and Ohta (1973)) are shown in black.

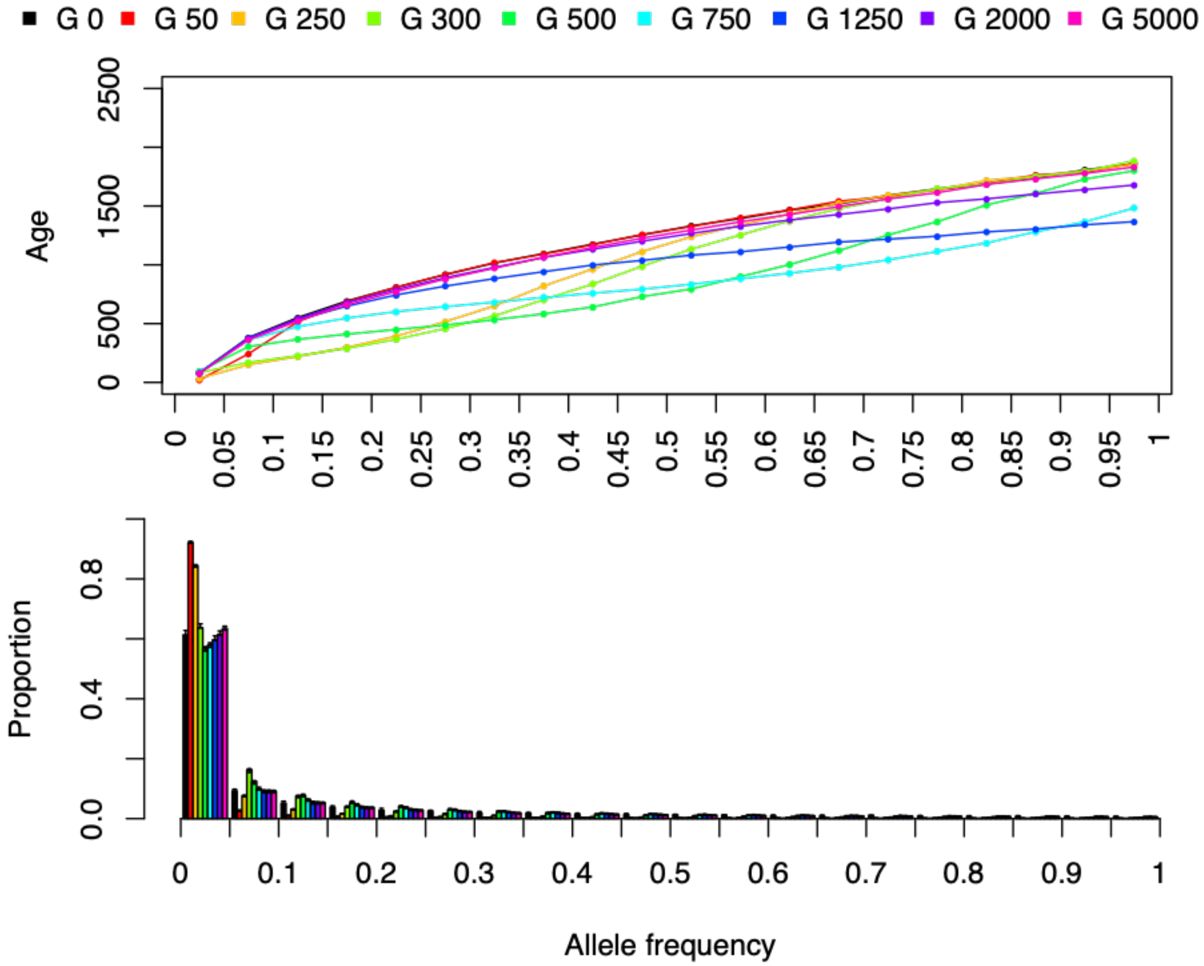


**Figure S3.** Mean age distribution and proportion of TEs at a given frequency in our TE burst model at the end of the burn in phase (G 0), generation 50 (G 50), 250 (G 250), 300 (G 300), 500 (G 500), 750 (G 750), 1250 (G 1250), 2000 (G 2000) and 5000 (G 5000). Top panel: mean TE age distribution. Bottom panel: TE site frequency spectrum (SFS).

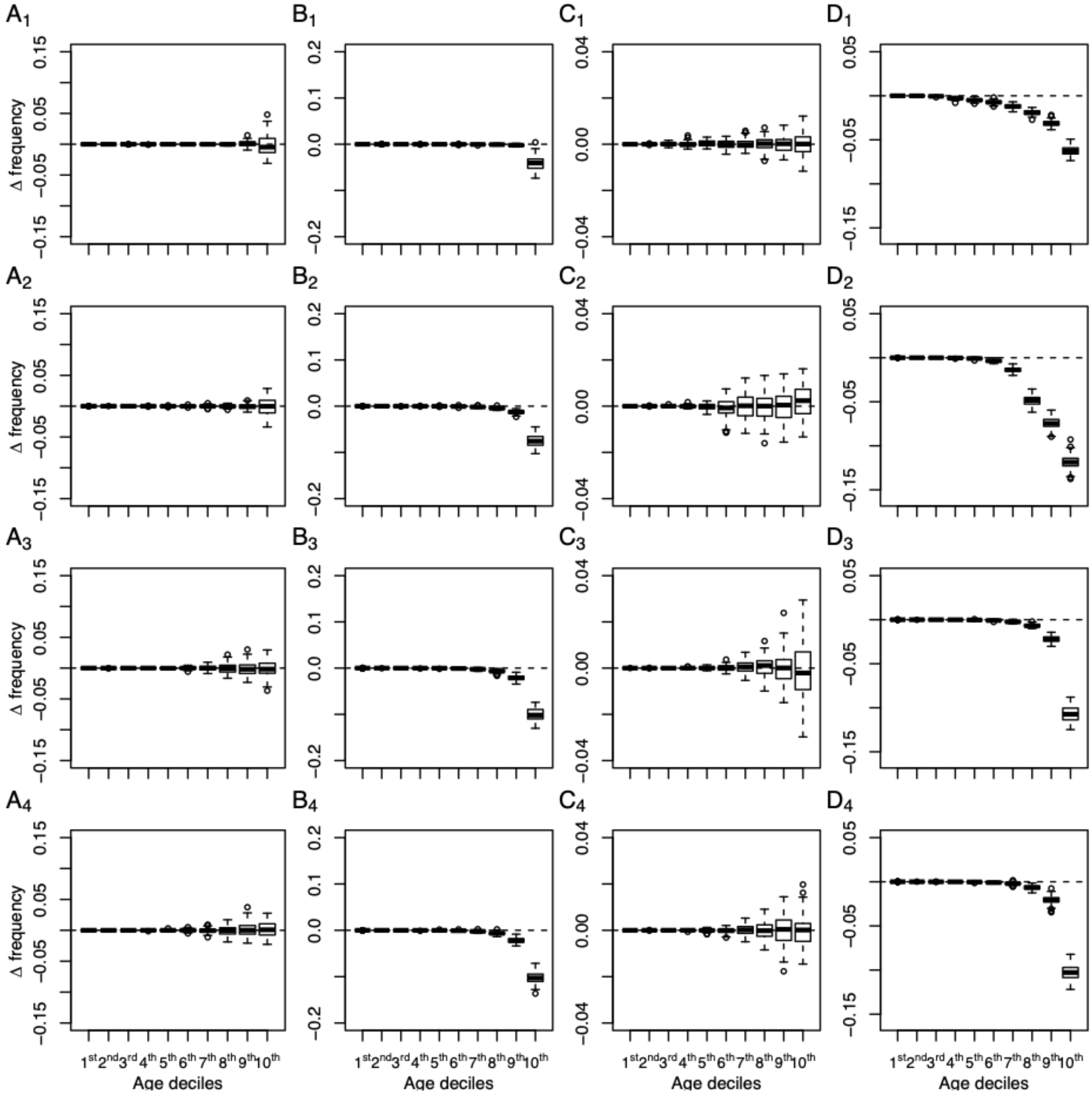


**Figure S4**. Age binned Δ frequency (mean TE frequency – mean SNP frequency) distributions observed in the different models. A. Observed Δ frequency between neutrally evolving TEs and SNPs under a bottleneck model. B. Observed Δ frequency between negatively selected TEs ($4N_es$ = -10) and neutrally evolving SNPs under a bottleneck model. C. Observed Δ frequency between neutrally evolving TEs and SNPs under a TE burst model. D. Observed Δ frequency between negatively selected TEs ($4N_es$ = -10) and neutrally evolving SNPs under a TE burst model. The four rows represent samples from generations 300, 500, 2000 and 5000 respectively.

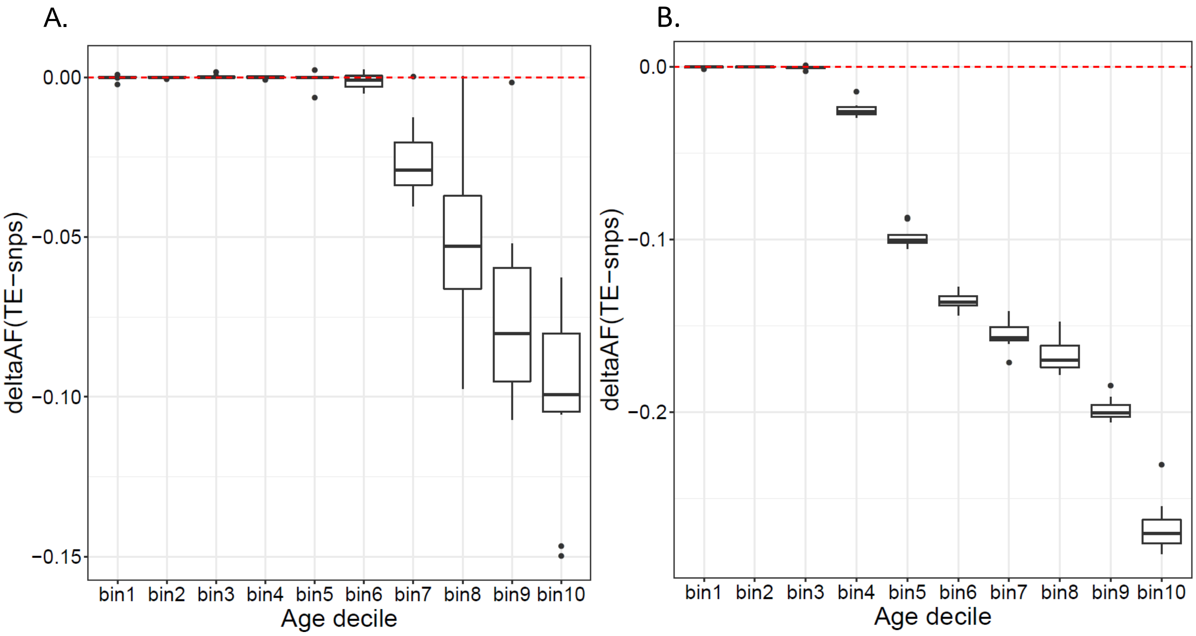


**Figure S5.** Age binned Δ frequency (mean TE frequency – mean SNP frequency) distributions after conditioning on empirical age estimates obtained from GEVA under A) a bottleneck model with negatively selected TEs ($4N_es$ = -10) and under B) a TE burst model with negatively selected TEs ($4N_es$ = -10). Note that singletons were not included in this analysis because of high inaccuracies in their age estimates.

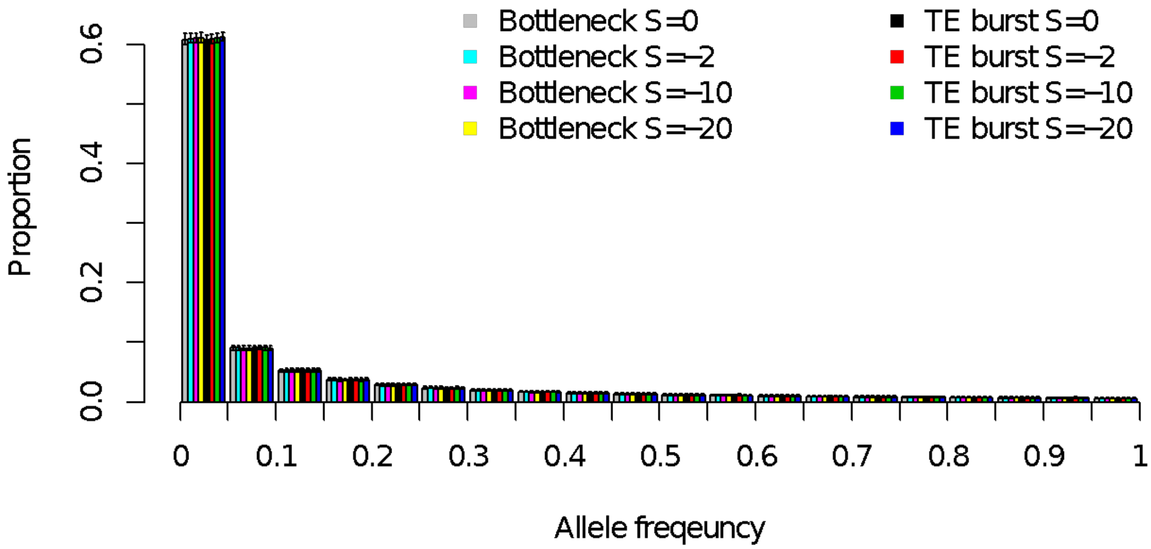


**Figure S6.** SNP SFS after the burn in phase in the different models.
